# Sex-specific loss of mitochondrial membrane integrity and mass in the auditory brainstem of a mouse model of Fragile X syndrome

**DOI:** 10.1101/2024.07.02.601649

**Authors:** Claire Caron, Elizabeth A. McCullagh, Giulia Bertolin

## Abstract

Sound sensitivity is one of the most common sensory complaints for people with autism spectrum disorders (ASDs). How and why sounds are perceived as overwhelming by affected people is unknown. To process sound information properly, the brain requires high activity and fast processing, as seen in areas like the medial nucleus of the trapezoid body (MNTB) of the auditory brainstem. Recent work has shown dysfunction in mitochondria, which are the primary source of energy in cells, in a genetic model of ASD, Fragile X syndrome (FXS). Whether mitochondrial functions are also altered in sound-processing neurons, has not been characterized yet. To address this question, we imaged the MNTB in a mouse model of FXS. We stained MNTB brain slices from wild-type and FXS mice with two mitochondrial markers, TOMM20 and PMPCB, located on the Outer Mitochondrial Membrane and in the matrix, respectively. These markers allow exploration of mitochondrial subcompartments. Our integrated imaging pipeline reveals significant sex-specific differences in the degree of mitochondrial length in FXS. Significant differences are also observable in the overall number of mitochondria in male FXS mice, however, colocalization analyses between TOMM20 and PMPCB reveal that the integrity of these compartments is most disrupted in female FXS mice. We highlight a quantitative fluorescence microscopy pipeline to monitor mitochondrial functions in the MNTB from control or FXS mice and provide four complementary readouts. Our approach paves the way to understanding how cellular mechanisms important to sound encoding are altered in ASDs.

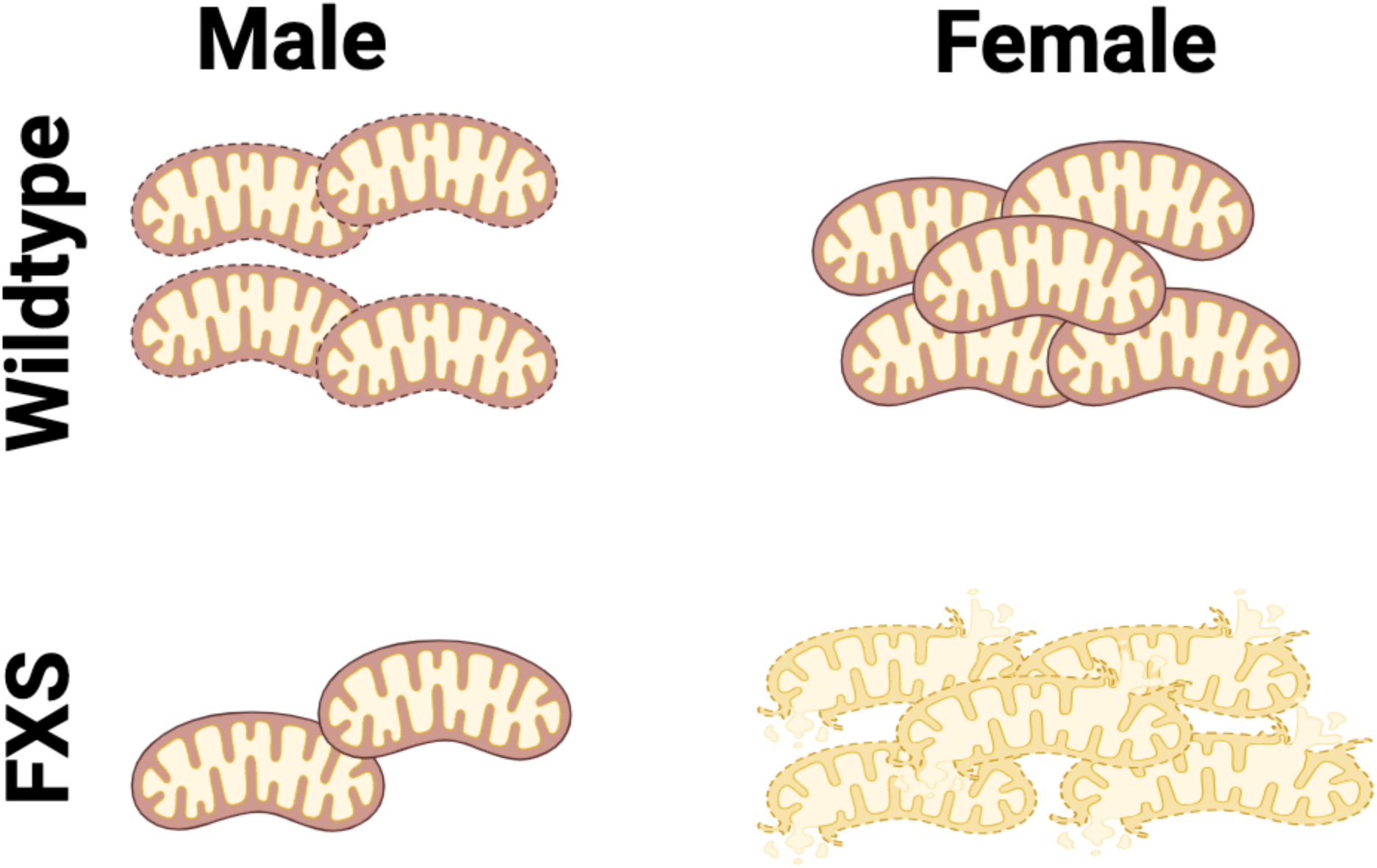

**Key points:**

- There are sex differences in mitochondrial structures in the MNTB.
- FXS female mice have disordered mitochondria that are also longer and less branched than wildtype females.
- Male FXS mice have fewer overall mitochondria compared to males.

## INTRODUCTION

Fragile X Syndrome (FXS) is the most common genetic cause of intellectual disability and autism spectrum disorder occurring in 1:4000-1:8000 individuals (Hagerman, 2008). FXS is caused by a CGG repeat expansion on the X chromosome in the *FMR1* gene resulting in reduced or no expression of the protein fragile X messenger ribonucleoprotein (FMRP). FMRP has important functions in the nervous system including regulating gene expression and its loss results in symptoms including cognitive impairments, sensory issues - including auditory sensitivity, anxiety, and overall hyperactivity. The role of FMRP in functioning of the nervous system has grown to not just include repressing specific mRNAs but binding to RNA, proteins, and channels resulting in the deficits seen in FXS (Richter & Zhao, 2021). Recent evidence also suggests that FMRP has roles in distinct functional pathways including energy metabolism, which may underlie multiple aspects of the disorder. Notably FXS patient fibroblasts showed donut shaped mitochondria, excessive mitochondrial proteins, and susceptibility to apoptotic cell death (Grandi *et al*., 2024). Other studies in a mouse model of FXS, *Fmr1* KO mice, showed an inner mitochondrial membrane leak, altered morphology, and impaired mitochondrial fusion in the hippocampus (Shen *et al*., 2019; Licznerski *et al*., 2020) and compartment specific plasticity which is lost in FXS (Bülow *et al*., 2021).

Mitochondria are multifunctional organelles which are essential for cell physiology and during stress response. They harbor the machinery devoted to energy production, the OxPhos complex, and they are key signaling hubs both internally and within cells. The wide range of metabolites produced by or transduced within mitochondria are important messengers not only for the different subcellular compartments, but also for intercellular communication (Picard & Shirihai, 2022). Interestingly, mitochondria as a whole are also a way for cells to communicate between them. Mitochondrial behaviors such as transport can be specialized in tissues within an organism to ensure specific cellular functions (Monzel *et al*., 2023). During typical mouse development, mitochondria are transferred between adjacent cells. This appears to be a multi-systemic feature occurring in several organs, including the brain (Marti Gutierrez *et al*., 2022). In the brain, the transfer of mitochondria from astrocytes to neurons was shown to be neuroprotective after stroke (Hayakawa *et al*., 2016). This strongly indicates that energetically-competent mitochondria are required at acute stress sites to overcome systemic failures.

Astrocyte functions were also shown to be impaired by a lower O2 availability and a lower mitochondrial energy production capacity in cultured astrocytes isolated from *Fmr1* KO mice (Vandenberg *et al*., 2022). While the overall mitochondrial ATP production levels were similar in male and female mice, this study showed that mitochondria from both wildtype and *Fmr1* KO male mice produced higher Reactive Oxygen Species (ROS) than those in their respective female counterparts. Therefore, it is important to consider sex as another key variable, potentially determining mitochondrial specificity in different brain cells (Gaignard *et al*., 2015; Ventura-Clapier *et al*., 2017).

The auditory brainstem, and particularly the medial nucleus of the trapezoid body (MNTB), is emerging as a model for neuroenergetics due to the high energy demands required of this brain area to be able to reliably generate action potentials up to 1000 Hz. Indeed energy demands, as measured by O2 consumption, of this brain area increase up to 600 Hz suggesting high energy demands that are dependent on temporal characteristics of stimulation (Palandt *et al*., 2024).

The auditory brainstem is required for sound localization processing, and the calyx of Held synapse which is the presynaptic terminal onto MNTB cells originating from globular bushy cells in the cochlear nucleus, is used both as a model synapse as well as critically for sound location computation. This is particularly relevant in regards to FXS, where auditory symptoms are common and some dysfunction may originate from early auditory brainstem function (McCullagh *et al*., 2020b).

A better understanding of the pathways involved in FXS is critical for developing therapies to remedy symptomology or rescue loss of FMRP. We hypothesize that mitochondrial functionality is critical to MNTB homeostasis and that the absence of FMRP leads to mitochondrial dysfunction. To this end, we explore the contribution of sex and the presence or absence of FMRP in mitochondria from MNTB neurons. In particular, we evaluate three complementary mitochondrial parameters including the morphology of the mitochondrial network, mitochondrial integrity, and mass. We uncover sex- and genotype-specific differences in mitochondrial length, integrity, and organelle mass, and we provide evidence that these parameters are correlated in MNTB neurons of FXS mice.

## METHODS

### Ethical Approval

All experiments complied with all applicable laws, National Institutes of Health guidelines, and were approved by the Oklahoma State University IACUC, approval number 20-07. Investigators understand the ethical principles under which the journal operates and that their work complies with the animal ethics checklist.

### Animals

Experiments were conducted in C57BL/6J (stock #000664, B6) wildtype background, hemizygous male and homozygous female *Fmr1* mutant mice (B6.129P2-*Fmr1^tm1Cgr^*/J stock #003025) obtained from the Jackson Laboratory (Bar Harbor, ME USA) and bred at Oklahoma State University (The Dutch-Belgian Fragile X Consorthium *et al*., 1994). Animals were generated for these experiments from stocks by both mixed and single genotype mating allowing for littermate controls as well as maintenance of breeding lines. Animals were maintained on individually ventilated racks in a room maintained on a 12:12 light:dark cycle with time on at 6AM and off at 6PM and fed ad libitum. Animals ranged in age from 77 – 113 days old (average ages per genotype 113 days old B6, 103 days old *Fmr1* KO). Numbers represent both sections from (number of animals), B6 males 106 images (3 animals), B6 females 77 images (3 animals), *Fmr1* KO males 115 images (5 animals), *Fmr1* KO females 49 images (3 animals). Variability in number of images and animals is due to blinding of the genotypes and subsequent genotyping of animals used in the study. Animals were genotyped using Transnetyx (Cordova, TN).

### Tissue Preparation

Tissue preparation was performed similar to previous work (McCullagh *et al*., 2017, 2022). Mice were anesthetized with pentobarbital (120 mg/kg body weight) and transcardially perfused with ice-cold phosphate buffered saline (PBS; 137 mM NaCl, 2.7 mM KCl, 1.76 mM KH2PO4, 10 mM Na2HPO4 Sigma-Aldrich, St. Louis, MO), followed by perfusion with 4% paraformaldehyde (PFA). Following perfusion, animals were decapitated, brainstems removed and post-fixed in 4% PFA overnight. Brainstems were then washed for ten minutes three times in PBS and covered in 4% agarose. Fixed brains were then sliced into 50 μm coronal sections with a Vibratome (Leica VT1000s, Nussloch, Germany) that included the medial nucleus of the trapezoid body (MNTB).

### Immunofluorescence

Once the brainstem was sliced into roughly 10 sections/animal, free-floating slices were blocked in antibody media (AB media: 0.1 M phosphate buffer (PB: 50 mM KH2PO4, 150 mM Na2HPO4), 150 mM NaCl, 3 mM Triton-X, 1% bovine serum albumin (BSA)) and 5% normal goat serum (NGS) for 30 minutes at room temperature on an orbital shaker. Following blocking, sections were incubated with an anti-PMPCB polyclonal antibody raised in rabbit (Proteintech Cat# 16064-1-AP, RRID:AB_2167122) and used at a 1:1000 dilution, with an anti-TOMM20 (Abcam Cat# ab56783, RRID: AB_945896) monoclonal antibody raised in mouse and used at a 1:5000 dilution, and 1% NGS in AB media without Triton-X overnight at 4°C on a rotating shaker. Slices were then washed (3 × 10 min) in PBS followed by secondary antibody goat anti-rabbit Alexa 647 (Invitrogen, Carlsbad, CA; H + L, cross adsorbed Cat#A21244) used at a 1:5000 dilution (RRID: AB_2535812), goat anti-mouse Alexa 488 (Invitrogen, Carlsbad, CA; H + L, highly cross adsorbed, Cat# A32723TR) used at a 1:5000 dilution (RRID: AB_2633275), and 1% NGS in AB media without Triton-X for one hour at room temperature on an orbital shaker. Slices were then washed in PB and mounted on glass slides with Fluoromount-G (SouthernBiotech, Cat.-No.: 0100-01, Birmingham, AL) and coverslipped (1.5 coverslip). At least 24 hours later, slides were taken to a Zeiss Airyscan at Oklahoma State to identify MNTB on slides which were circled on the reverse side of the slide for identification in Rennes. Animal identification was then blinded and slides were shipped cold overnight for imaging in Rennes.

### Antibody characterization

The anti-TOMM20 primary antibody was used for the detection of the outer mitochondrial membrane, while the anti-PMPCB primary antibody was used to label the inner mitochondrial membrane/matrix compartment. To visualize these primary antibodies, two complementary fluorescent-conjugated secondaries were used. The goat anti-mouse Alexa 488 was used to detect the anti-TOMM20 primary antibody, and the goat anti-rabbit Alexa 647 was used to detect the anti-PMPCB primary antibody. The anti-PMPCB antibody (Proteintech Cat# 16064-1-AP, RRID:AB_2167122) is a polyclonal antibody that is specific to the PMPCB fusion protein Ag8937 which corresponds to a 1-349 amino acid sequence encoded by BC010398. This antibody was previously validated for *PMPCB* knockdown efficiency (Greene *et al*., 2012; Bertolin *et al*., 2018). The anti-TOMM20 primary antibody is a mouse monoclonal antibody specific to the human TOMM20 at amino acids 1-145. TOMM20 staining was validated through its co-localization with PMPCB (but not overlap) in addition to consistent staining pattern with other citations using the same antibody in the brain (Pease-Raissi *et al*., 2017).

### Imaging

All slides were imaged blinded to genotypes. Multicolor images were acquired as individual optical sections in *xy*, with a Leica SP5 inverted confocal microscope (Leica) driven by the Leica Acquisition Suite (LAS) software, and a 63X (N.A. 1.4) oil immersion objective. The excitation/emission wavelengths for Alexa 488 were 488 and 525/50 nm respectively; for Alexa 647 were 633 and 650/20 nm, respectively. The 488 nm laser power was set at 40%, and the 633 nm laser power was set at 20% throughout conditions. For visualization purposes only, representative images were deconvolved using the Fiji DeconvolutionLab plugin and the Tikhonov-Miller algorithm, with built-in settings used throughout conditions. Brightness and contrast were adjusted and applied to the entire image.

### Quantification of Images

Raw images were used for quantification throughout the manuscript. Fluorescence colocalization (Mander’s M1 and M2 coefficients) between TOMM20 and PMPCB was calculated with the JaCoP plugin (Bolte & Cordelieres, 2006) of the Fiji software (NIH), and the built-in automatic threshold mask was applied to the confocal images. Mitochondrial length and branching were calculated on the PMPCB-specific staining, using previously validated procedures to extract Aspect Ratio and Form Factor coefficients as indicators of mitochondrial morphology (Koopman *et al*., 2006; Buhlman *et al*., 2014). Briefly, Aspect Ratio is the ratio between the major and minor axes of each mitochondrial object, whereas Form Factor is calculated by performing (1/Circularity). For each mitochondrial object, Aspect Ratio and Circularity were extracted thanks to the Fiji feature “Shape descriptors”, and then Form Factor was calculated for each Circularity value. This procedure also provides the overall number of PMPCB-positive mitochondrial objects, which was then used as a readout of mitochondrial mass.

### Statistical Analysis

Ordinary two-way ANOVA tests were performed with GraphPad Prism v. 10, and used to compare the sex and genotype for each mitochondrial parameter tested. All tests were performed after testing data for normality. For all tests, alpha was 0.05. The results of each test were plotted as violin plots showing the entire data distribution, and containing *n*=90 MNTB neurons from three WT males, 60 MNTB neurons from three WT females, 120 MNTB neurons from five *Fmr1* KO males and 90 MNTB neurons from three *Fmr1* KO female mice. Significant and non-significant comparisons, together with their respective *P* values, are indicated in each corresponding Figure panel. Factor analysis of mixed data (FAMD) was performed with Rstudio v.2024.04.0-735, and using the functions associated with the *FactoMineR* and *Factoextra* libraries, including the Fviz functions. Genotype and sex were used as qualitative variables. Mander’s M2 coefficient and mitochondrial numbers were used as quantitative variables, with *n*=90 MNTB neurons from three WT males, 60 MNTB neurons from three WT females, 120 MNTB neurons from five *Fmr1* KO males and 90 MNTB neurons from three *Fmr1* KO female mice. The correlation circle was built using quantitative variables only, while the FAMD maps were built using quantitative and qualitative variables. Quantitative and qualitative variables were normalized during the analysis in order to balance the influence of each set of variables

## RESULTS

### 1. Mitochondrial morphology of MNTB cells is mildly affected in FXS mice

Previous reports showed altered mitochondrial morphologies in FXS models. In fibroblasts generated from FXS patients, mitochondria display a donut-like shape and have an increased number of cristae when compared to controls (Grandi *et al*., 2024). A decreased ER-mito distance is also observable in the hippocampus of *Fmr1* KO mice (Geng *et al*., 2023), and it is known that mitochondrial morphology depends on the proximity of the organelles with the Endoplasmic Reticulum (ER) (Marchi *et al*., 2014).

Since FXS patients and mouse models show alterations to their auditory system (McCullagh *et al*., 2020b), we hypothesized that mitochondrial morphology alterations are also present in *Fmr1* KO mice MNTB neurons. To this end, we first co-immunostained MNTB sections of wildtype (B6) and *Fmr1* KO mice with the mitochondrial outer membrane marker (OMM) TOMM20, and with the matrix marker PMPCB. Second, MNTB neurons from male and female wildtype and KO mice were imaged with confocal microscopy (Fig. 1). Then, we used the PMPCB-specific signal to calculate mitochondrial aspect ratio and form factor (Fig. 2). These two parameters report on organelle length and branching, respectively (Koopman *et al*., 2006). We observed that mitochondrial length in MNTB neurons is similar between male and female wildtype mice, and that male *Fmr1* KO mice were no longer than their wildtype counterparts (Fig. 2A-B, *P*=0.055). However, a significant difference was detected between female wildtype and *Fmr1* KO mice, indicating an increased mitochondrial length in FXS females (Fig. 2B, *P*<0.0001). Mitochondria showed a higher degree of mitochondrial branching in MNTB neurons from female wildtype mice when compared to their male counterparts (Fig. 2A, C, *P*=0.002). While no difference between wildtype and *Fmr1* KO males was detected (*P*=0.383), mitochondria from female wildtype mice showed a higher degree of mitochondrial branching than their male counterparts (Fig. 2C, *P*=0.022). In female *Fmr1* KO mice, mitochondrial branching was similar to that observed in males (Fig. 2C, *P*=0.337).

**Figure 1:**
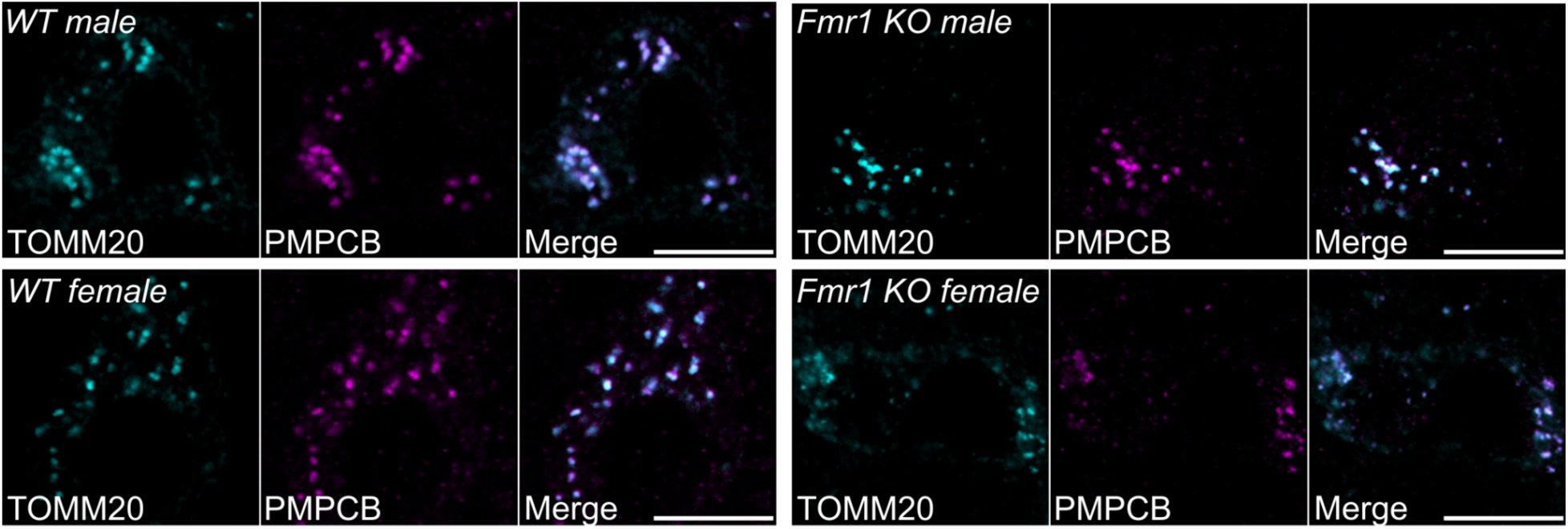
Representative fluorescent micrographs of MNTB neurons co-stained with antibodies against the OMM marker TOMM20 (pseudocolor cyan) and the IMM/matrix marker PMPCB (pseudocolor magenta), in both male and female wildtype (WT) or *Fmr1* KO mice. TOMM20 and PMPCB were merged to show marker colocalization. Images were acquired as individual optical sections in xy, with a Leica SP5 confocal microscope and a X63 (N.A. 1.4) oil objective. The representative images were deconvolved using the Fiji DeconvolutionLab plugin and the Tikhonov-Miller algorithm for visualization purposes only. Brightness and contrast were adjusted for visualization purposes only. Scale bar: 10 µm.

**Figure 2:**
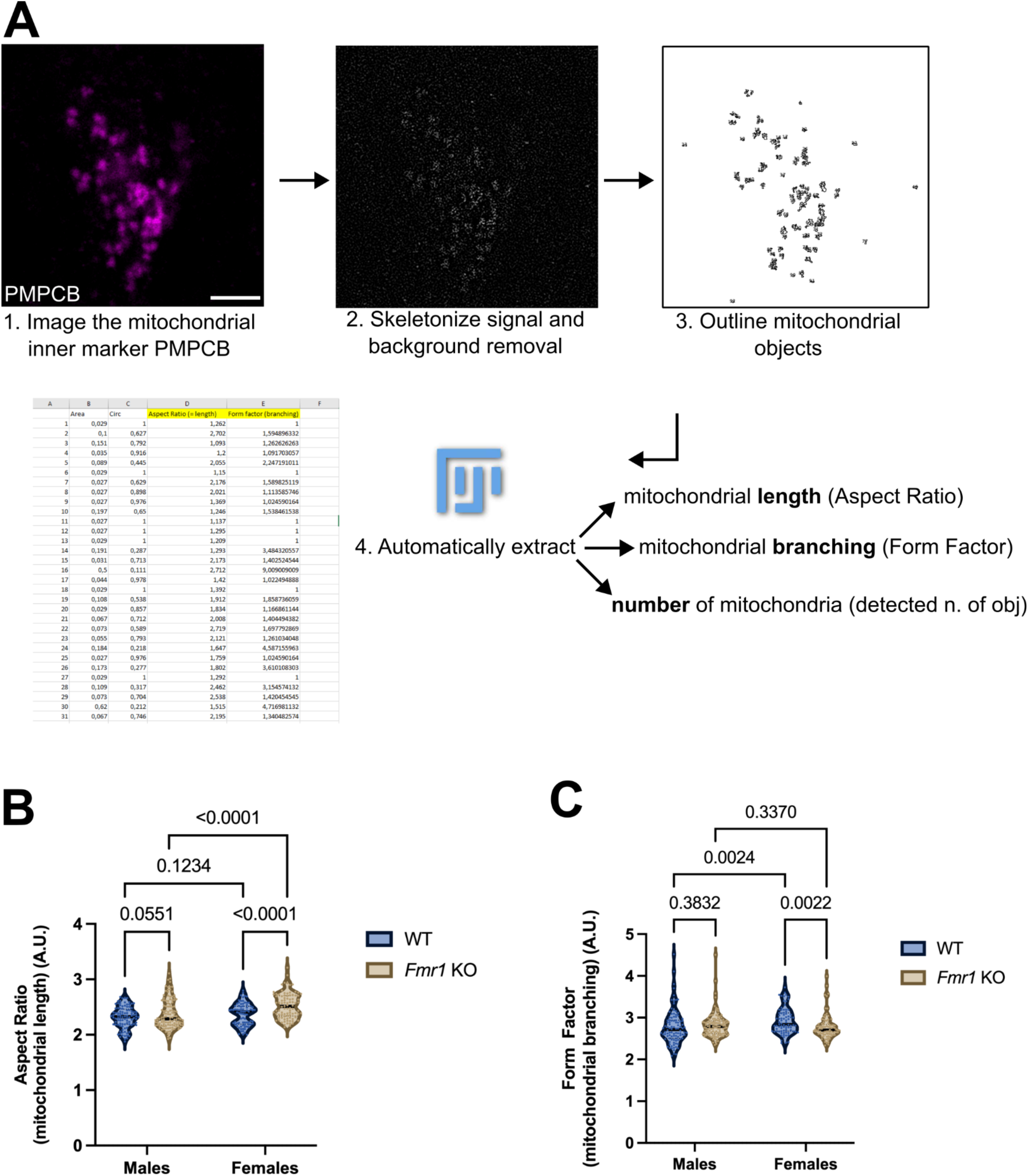
(**A**) Fiji-based image analysis pipeline to automatically extract mitochondrial length, branching and the overall number of mitochondria from confocal images. Once PMPCB-positive, xy individual optical sections were obtained as in Fig.1 (representative micrograph, pseudocolor magenta) (1), the signal is skeletonized (2) to obtain mitochondrial objects (3). Each mitochondrial-positive object has a size of ≥ 15 px. All objects smaller than 15 px were considered as background and discarded. For every object, the Aspect Ratio and the Form Factor were calculated following (Koopman *et al*., 2006; Buhlman *et al*., 2014). The overall number of PMPCB-positive objects of ≥ 15 px was also extracted. (**B**) Violin plots of Aspect Ratio and (**C**) Form Factor analyses in male and female mice, either wildtype (WT) or KO for *Fmr1*. All analyses were performed as in (**A**). *n*=90 MNTB neurons from three WT males, 60 MNTB neurons from three WT females, 120 MNTB neurons from five *Fmr1* KO males and 90 MNTB neurons from three *Fmr1* KO female mice. Data extends from min to max. The median is shown as a bar and individual *n* values are shown as dots in each condition. Significant and non-significant comparisons, together with their respective *P* values, are indicated. A.U.: arbitrary units.

Overall, confocal imaging of MNTB neurons shows sex-specific differences in mitochondrial length and interconnectivity, which is regulated by the presence or absence of *Fmr1*. We observed that mitochondria are significantly longer in female *Fmr1* KO mice than in any other condition and regardless of sex or genotype. In female *Fmr1* KO mice, mitochondria are also less interconnected than in wildtype females. Lastly, we observe that mitochondria are more interconnected in MNTB neurons from wildtype females than in those from males, thereby highlighting a sex-specific difference.

### 2. Mitochondrial membrane integrity in MNTB cells is sex-specific, and it is decreased in female FXS mice

In humans, FXS affects males roughly two times more than females both in prevalence and clinical manifestation (Rinehart *et al*., 2011). Both mitochondrial function and auditory brainstem physiology show sex-specific differences in FXS astrocyte culture and mouse models (Chawla & McCullagh, 2022; Vandenberg *et al*., 2022). Indeed, mitochondrial structure and function have been linked to sex differences in several pathologies (Ventura-Clapier *et al*., 2017) and it is increasingly accepted that mitochondria are sexually dimorphic in metabolism and cell death (Demarest & McCarthy, 2015). The integrity of the inner and outer mitochondrial membrane (IMM and OMM respectively) are critical to the function of mitochondria. When the integrity of the membranes is compromised, there are alterations to the membrane potential with depletion of ATP generation capacity and energy compromise (Cowan *et al*., 2019; Han *et al*., 2020). A loss of membrane integrity is also key to initiate mitochondrial turnover pathways by mitophagy (Chan *et al*., 2011; Pickrell & Youle, 2015; Wang *et al*., 2023). To address whether the ratio of the IMM and OMM were compromised in the MNTB of FXS, we calculated both the Mander’s M1 (PMPCB-IMM to TOMM20-OMM) and M2 (TOMM20-OMM to PMPCB-IMM) colocalization coefficients using the stained sections imaged above in male and female FXS mouse MNTB (Fig. 3A). We found no significant difference between male genotypes in the M1 and M2 coefficients (*p*=0.178 and *p*=0.669, respectively). In contrast, female FXS mice showed significant reductions in M1 and M2 coefficients compared to wildtype females, suggesting a decreased mitochondrial membrane integrity (Fig. 3B, C, *p*<0.0001 for both M1 and M2). Interestingly, there were also sex-specific differences in wildtype M1 and M2 coefficients. We observed higher M1 and M2 coefficients in wildtype females compared to wildtype males (*P*=0.001 and *P*=0.003, respectively). Lastly, FXS males also showed higher M1 (*P*=0.0001) and M2 coefficients (*P*<0.0001) compared to FXS females (Fig. 3B-C).

**Figure 3:**
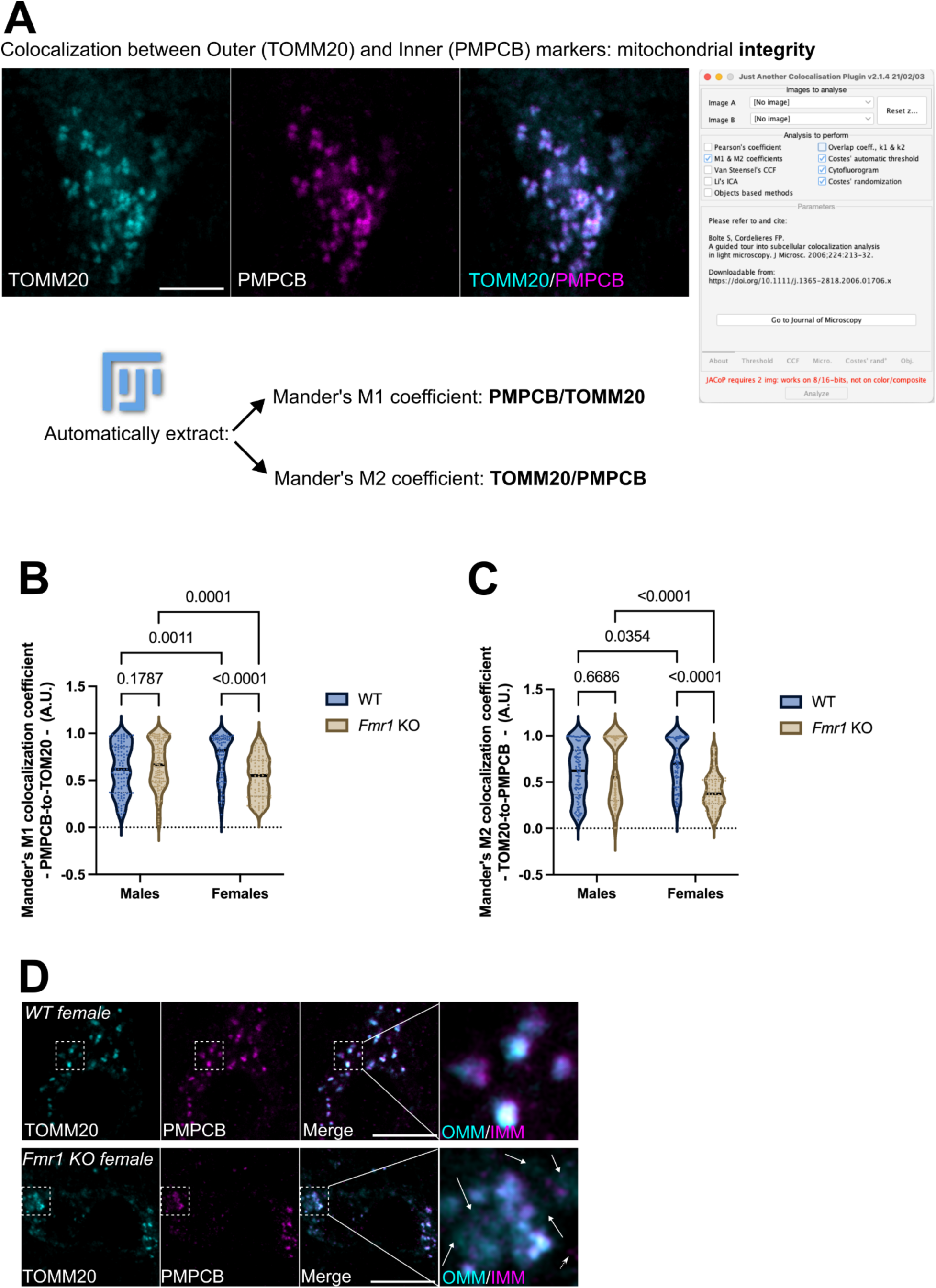
(**A**) Representative fluorescent micrographs and Fiji-based image analysis pipeline to automatically extract mitochondrial integrity from confocal images using the JACoP plugin (Bolte & Cordelieres, 2006). Once TOMM20- (pseudocolor cyan) and PMPCB- (pseudocolor magenta) positive xy individual optical sections were obtained as in Fig. 1, Mander’s M1 and M2 coefficients were calculated using the JACoP plugin. The performance of these calculations was controlled with Costes’ automatic threshold, randomization and with a cytofluorogram directly with JACoP built-in solutions, and this for every pair of images analyzed. (**B**) Violin plots of Mander’s M1 and (**C**) of Mander’s M2 coefficients in male and female mice, either wildtype (WT) or KO for *Fmr1*. All analyses were performed as in (**A**). *n*=90 MNTB neurons from three WT males, 60 MNTB neurons from three WT females, 120 MNTB neurons from five *Fmr1* KO males and 90 MNTB neurons from three *Fmr1* KO female mice. Data extends from min to max. The median is shown as a bar and individual *n* values are shown as dots in each condition. Values span from 0 to 1, where 0 indicates no colocalization and 1 indicates perfect colocalization. Significant and non-significant comparisons, together with their respective *p* values, are indicated. (**D**) Representative fluorescent micrographs of MNTB neurons co-stained with antibodies against the OMM marker TOMM20 (pseudocolor cyan) and the IMM/matrix marker PMPCB (pseudocolor magenta), in wildtype (WT) or *Fmr1* KO female mice. TOMM20 and PMPCB were merged to show marker colocalization. Dotted area: magnified Region Of Interest (ROI) to illustrate membrane integrity loss in *Fmr1* KO mice compared to wild-type. Arrows show TOMM20-positive, but PMPCB-negative mitochondria. The dotted arrow shows PMPCB-positive, but TOMM20-negative mitochondria. Images were acquired as individual optical sections in xy, with a Leica SP5 confocal microscope and a X63 (N.A. 1.4) oil objective. The representative images were deconvolved using the Fiji DeconvolutionLab plugin and the Tikhonov-Miller algorithm for visualization purposes only. Brightness and contrast were adjusted for visualization purposes only. Scale bar: 10 µm. A.U.: arbitrary units.

In summary, FXS females show the most severe membrane phenotype, with decreased integrity in both inner and outer membrane ratios. In MNTB neurons, FMRP loss leads to mitochondria with either OMM- or matrix-only markers (Fig. 3D). This might lead to decreased ATP production in the MNTB of FXS females. We also highlight sex-specific deficits in mitochondrial membranes. Wildtype females have increased M1 and M2 compared to males, thereby indicating a potential sexual dimorphism of mitochondrial membrane integrity in the MNTB.

### 3. Male FXS mice show mitochondrial mass loss in MNTB

Prior data obtained from fibroblasts of FXS patients indicated that there is either no overall loss of mitochondrial proteins (Geng *et al*., 2023), or an increase of specific proteins related to mitochondrial respiration such as the ATPase inhibitor IF1, cyclophilin D (CyPD), or the ATP synthase OSCP subunit (Grandi *et al*., 2024). However, MNTB mitochondria showed a loss of membrane integrity in FXS female mice (Fig. 3B-D). We reasoned that this could be the result of an overall loss of organelle mass under these conditions. In addition, work in cell models also showed a link between FMRP and the efficient clearance of defective mitochondria by mitophagy (Geng *et al*., 2023).

To address this possibility, we calculated the number of PMPCB-positive organelles, which reflects the mitochondrial mass, across sexes and genotypes using confocal microscopy and image analysis (Fig. 2A). Our analyses revealed no difference in the number of mitochondrial objects between wildtype and *Fmr1* KO female mice (Fig. 1, 4A, *p*=0.7252). Surprisingly, similar analyses revealed a significant loss of PMPCB-positive objects in *Fmr1* KO males compared to their wildtype counterparts (Fig. 4A, *p*≤0.0001).

**Figure 4:**
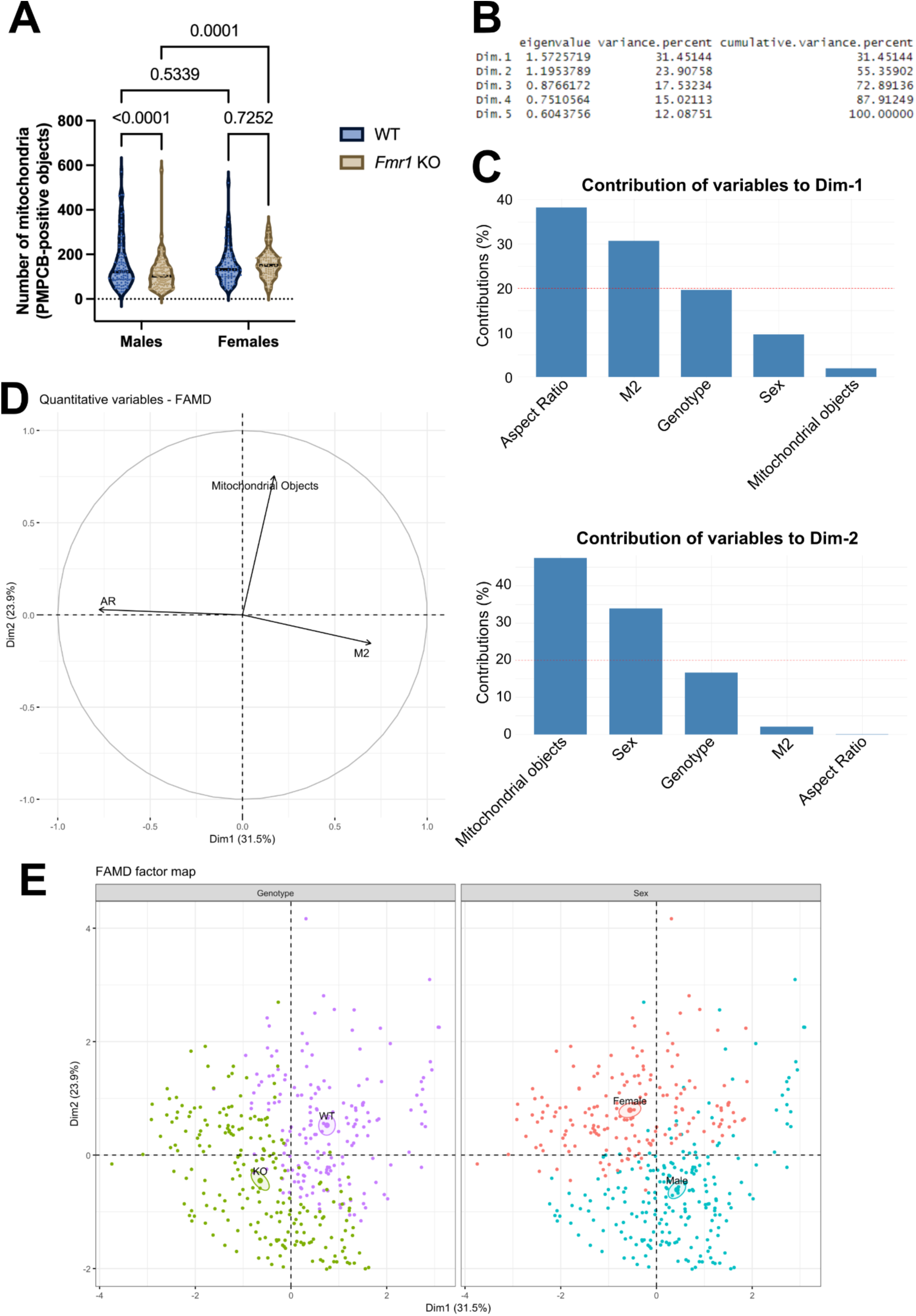
(**A**) Violin plots of the number of PMPCB-positive mitochondrial objects in male and female mice, either wildtype (WT) or KO for *Fmr1*. All analyses were performed as in Fig. 2A. *n*=90 MNTB neurons from three WT males, 60 MNTB neurons from three WT females, 120 MNTB neurons from five *Fmr1* KO males and 90 MNTB neurons from three *Fmr1* KO female mice. Data extends from min to max. The median is shown as a bar and individual *n* values are shown as dots in each condition. Significant and non-significant comparisons, together with their respective *P* values, are indicated. A.U.: arbitrary units. (**B**) Raw and percentage of eigenvalue variance, and percentage of cumulative variance of data dimensions in our datasets. In the cumulative variance, values add up to reach the entire (100%) dataset. (**C**) Contribution of each indicated variable to the first data dimension (in percentage, upper graph) or to the second data dimension (in percentage, lower graph). The red dashed line indicates the expected average value. (**D**) Correlation plot between Aspect Ratio, the number of mitochondrial objects and the Mander’s M2 coefficient across the first and second data dimensions. (**E**) FAMD factor score plots showing the genotype- and sex-dependent clustering of the entire dataset across the first and second data dimensions. Individual *n* values as in (**A**) are shown as dots. Pseudocolors green and purple were used to represent the genotype, while pseudocolors blue and red were used to represent the sex.

So far, we observed that mitochondrial number, but not integrity, was lowered in *Fmr1* KO males, and that mitochondria were longer and less intact in *Fmr1* KO females. Therefore, we asked whether organelle length, integrity and number may be correlated in MNTB neurons across sex and genotype. To this end, we performed Factor Analysis of Mixed Data (FAMD), a method suitable to analyze mixed datasets. Indeed, FAMD allows the analysis of quantitative variables as a Principal Component Analysis (PCA), and of qualitative values as a Multiple Factor Analysis (MFA) (Pagès, 2004). In our analyses, the Aspect Ratio, Mander’s M2 coefficient, and mitochondrial number were used as quantitative variables, while sex and genotype were used as qualitative variables. First, we determined that our dataset can be described by five data dimensions, and that the first and the second data dimensions together account for 55% of all the variables needed to illustrate the entire dataset (Fig. 4B). We then evaluated the contribution of each quantitative and qualitative variable to the main two data dimensions. We observed that the Aspect Ratio, the Mander’s M2 coefficient and the genotype largely contribute to the first data dimension. In contrast, mitochondrial objects and the sex are the main variables representing the second data dimension (Fig. 4C).

We then explored the correlation between the three quantitative variables - mitochondrial length, integrity and number - regardless of sex and genotype. We found that the Mander’s M2 coefficient and the Aspect Ratio, both contributing to the first data dimension, are anti-correlated (Fig. 4D). This corroborates previous data shown in Fig. 2C and 3C and shows that the longer the organelles, the lower the integrity. Furthermore, the Mander’s M2 coefficient and mitochondrial objects, which are contributing largely to the second data dimension, are positively correlated (Fig. 4D). Lastly, we evaluated how genotype (WT vs *Fmr1* KO) and sex (male vs female) are distributed in the two data dimensions, for each MNTB neuron imaged within the dataset. This representation integrates the two qualitative variables across the two main dimensions which are mainly shaped by quantitative variables (Fig. 4C). According to the FAMD factor maps, we observed that MNTB neurons can be segregated into two main groups either in terms of genotype (left score plot) or sex (right score plot) (Fig. 4E). Overlap between the two clusters was limited. In the left score plot, we observed that mitochondria from KO MNTB neurons show a positive correlation between Aspect Ratio (Dim. 1) and the number of mitochondrial objects (Dim. 2), while showing an anti-correlation between Aspect Ratio and the Mander’s M2 coefficient (Dim. 1) (Fig. 4E). When looking at sex specific effects in the right score plot, we observe that the MNTB neurons with longer, more abundant and less intact mitochondria are those from female mice. Conversely, mitochondria from males have a low amount of mitochondrial objects, a low Aspect Ratio and a high proportion of intact mitochondria.

## DISCUSSION

We provide evidence that mitochondria are altered in MNTB neurons of mice affected with FXS. In FXS mice, mitochondrial alterations occur at at least three different levels including a change in the morphology of the mitochondrial network, membrane integrity, and organelle mass. These findings have significant implications on the function of auditory mitochondria which could explain some of the auditory specific phenotypes seen in mouse models and people with FXS due to the high energetic demands of this region. Specifically, we show sex differences in mitochondrial structures with FXS females having longer mitochondria with less integrity that are more abundant than in FXS male mice. In contrast, males exhibit more mitochondria than wildtype mice, potentially suggesting that different pathways may be involved in FXS phenotypes between the sexes.

As previously described, mitochondrial respiration and ROS production in astrocytes of FXS mice were reported to be sex-specific (Vandenberg *et al*., 2022). However, this study was performed in isolated primary astrocytes which were then cultured, permeabilized and processed for *in vitro* analyses. First, it is worth considering that astrocytes and MNTB neurons have different physiology and may show intrinsic variability in mitochondrial functions - including metabolism. Second, isolating and culturing astrocytes may result in these cells adapting their functions *ex vivo*. As a consequence, this may explain why the majority of sex-specific differences were overall modest, and were only present in hypoxic conditions. The sex-specific differences observed in MNTB neurons not only expand the original findings generated in astrocytes to an alternative neuronal population, but also show that significant mitochondrial morphology, integrity and mass differences exist at the tissue level. Lastly, a recent study performed in mice quantified the maternal transmission of an active *Fmr1*-KO allele to be higher than the paternal transmission (Szelenyi *et al*., 2024). This finding may explain why the mitochondrial phenotype is stronger in MNTB neurons from female KO mice than what we observe in males. This also raises the interesting possibility that mitochondrial signaling pathways may be differentially regulated in males and females. In this light, the presence of local chromosome X activation/inactivation mosaicism correlates with the prevalence of specific symptoms (Szelenyi *et al*., 2024). These differences could also be translated at the subcellular level, where the overall decrease in the number of mitochondria observed in males may result from a loss of membrane integrity in females. Although future studies are required to corroborate this possibility, this may pave the way to the integration of sex specificity and the mitochondrial phenotypic variability in therapeutic solutions against FXS.

Despite the importance of the auditory brainstem in terms of energetic demands, for example being able to generate action potentials up to 1000 Hertz (Kopp–Scheinpflug *et al*., 2003), few studies have focused on mitochondrial function in this region (Trattner *et al*., 2013; Lujan *et al*., 2016, 2021; Lucas *et al*., 2018; Palandt *et al*., 2024). While other studies included both male and female animals, due to the constraints of *in vitro* preparations, or developmental questions, they were all performed at young ages <postnatal day 40 where it is unexpected for there to be sex differences and/or it is difficult to discriminate the sexes (Trattner *et al*., 2013; Lujan *et al*., 2016, 2021; Lucas *et al*., 2018; Palandt *et al*., 2024). To our knowledge, this is the first study to characterize sex differences in auditory brainstem mitochondria, which has important implications for sexes differences in sound information independent of FXS. For example, mitochondria of wildtype males were significantly less intact, had fewer branches, and were shorter than wildtype females suggesting underlying ATP production differences that, if possible, should be explored in adult animals. Another interesting area of future research would be what drives sex differences specifically in this area, if there are cyclical differences in females suggesting estrus specific effects, or if there are other hormonal contributions to mitochondrial function in this area, which may underlie differences in sound localization in the sexes (McCullagh *et al*., 2020a).

Mitochondrial ultrastructure, morphology, respiratory capacity and sensitivity to apoptosis were recently evaluated in cultured fibroblasts from FXS patients (Grandi *et al*., 2024). These complementary parameters were used to define the direct impact of FXS-related mutations on mitochondrial functions. In the present study, an integrated imaging pipeline allowed quantification of mitochondrial phenotypes directly on a tissue relevant for the disorder. This pipeline, which is entirely based on open-source software, could easily be implemented to explore alternative mitochondrial markers. This could allow us to image additional organelle functions, or the regulation of mitochondrial signaling pathways. The imaging pipeline could also be adapted to alternative brain areas, or to measure mitochondrial functions in the auditory brainstem of alternative model organisms.

The auditory brainstem provides a unique model to test for mitochondrial changes in an area of the brain that requires some of the most energetic synapses in order to encode sound information. In addition there are advantages to using anatomical approaches in adult animals to characterize mitochondrial phenotypes such as the ability to correlate changes with auditory phenotypes in FXS such as altered auditory brainstem responses (Chawla & McCullagh, 2022) or *in vitro* physiology (Brown *et al*., 2010; Strumbos *et al*., 2010; Curry *et al*., 2018; El-Hassar *et al*., 2019; Lu, 2019) that is typically limited to younger developmental ages. Understanding adult phenotypes is an important step towards testing for efficacy of drug interventions that might target mitochondria in patients that already have FXS. In addition, while powerful for understanding mechanisms, it can be difficult to connect fibroblast or culture studies with biological relevant systems, like auditory processing.

The integrated imaging pipeline presented here is currently optimized for images acquired in two dimensions. Therefore, future improvements in the pipeline should determine whether the mitochondrial phenotypes observed in this study are limited to the axial dimension or if they extend to the lateral dimension as well. Additionally, the pipeline needs to be calibrated to define the minimal size of a mitochondrial-specific object. This parameter is mandatory for executing the entire pipeline, as it allows to distinguish the organelles from the background. This calibration is rather straightforward when imaging abundant mitochondrial proteins such as PMPCB or TOMM20.

Nevertheless, alternative mitochondrial markers may have a higher signal-to-noise ratio, or may be unevenly distributed at the sub-mitochondrial level. This could hamper the calibration of the pipeline, and may give rise to different results than those reported here.

One of the biggest limitations to the current study is that it is anatomical only and does not necessarily extend to functional changes that might be occurring with mitochondria in the auditory brainstem. Future studies should extend mitochondrial changes to functional auditory brainstem outputs such as respirometry or *in vivo* markers for mitochondrial function. One limitation to the auditory brainstem is that these cells can be difficult to culture in monolayer and maintain their specific characteristics, therefore organotypical sections are required which can be more difficult to maintain and are low-throughput. In particular, understanding of the physiological consequences of sex differences especially regarding physiological phenotypes of longer, less intact female FXS mitochondria is an area of needed future research.

### Future directions

A previous report provided a connection between mitochondrial mass loss and Parkin/PINK1-related mitophagy in FXS models (Geng *et al*., 2023). Our study shows a loss of mitochondria in MNTB neurons from male FXS mice. Therefore, future analyses in this neuronal population will demonstrate whether Parkin/PINK1-related mitophagy triggers the observed mitochondrial mass loss, or if alternative mitophagy pathways are implicated. Complementary information could also be obtained with ultrastructural data, where the morphology and the integrity of individual mitochondria is estimated with electron microscopy-based analyses. This approach would also allow to directly compare the ultrastructural changes previously observed in fibroblasts (Grandi *et al*., 2024) and in MNTB neurons.

Furthermore, determining the mitochondrial energetic capacity in MNTB neurons from FXS mice remains pivotal. This would provide information on how FMRP may impact energy metabolism in a neuronal population with high energy demands (Palandt *et al*., 2024). A previous study highlighted the role of FMRP in regulating the number of ER-mitochondria contact sites (ERMCSs) (Geng *et al*., 2023). FMRP-dependent ERMCSs were shown to be key not only for energy metabolism, but also to maintain synaptic structure and function in a *Drosophila* model of FXS through Ca^2+^ dynamics.

Therefore, our results pave the way to functionally correlate mitochondrial mass loss, the pathways regulating mitochondrial energetic capacity, and the molecular actors orchestrating these functions in MNTB neurons.

Presynaptic release requires a lot of energy to maintain neurotransmission, particularly at high frequency stimulation rates. Surprisingly, the calyx of Held presynaptic synapse uses glycolysis to maintain neurotransmission at baseline and low-frequency stimulation (Lujan *et al*., 2016). While we do not discriminate in our study between pre and post synaptic mitochondria, it is expected based on energetic demands that the majority of our measured mitochondria are located presynaptically.

However, due to different energy demands between the pre and post synapse, it would be interesting to differentiate between mitochondria at either side of the synapse to show if there are location-specific alterations in mitochondrial form and function. Higher-resolution imaging such as electron microscopy would be necessary and can provide finer detail regarding mitochondrial form as well to corroborate our results. Differences in energy demands at rest, low, and high frequency in the presynaptic terminal of FXS mice would also give important insights into the pathways involved, whether glycolytic or due to mitochondrial respiration alterations.

In conclusion, we here show that mitochondrial length, mass, and integrity are correlated in MNTB neurons across genotype and sex. With our integrated analysis pipeline, we demonstrate that *Fmr1* KO males have shorter, less abundant but more intact mitochondria. In contrast, *Fmr1* KO females have longer, less intact but an overall equal amount of mitochondria to their wildtype counterparts. Our data not only show that FMRP is key for the maintenance of mitochondrial physiology in MNTB neurons, but also highlight sex-specific differences that may arise in this specific neuronal population.

## ADDITIONAL INFORMATION

### Data access

Data is available upon request.

### Author contributions

Animal experiments were performed at Oklahoma State University, including housing and care of animals, and immunofluorescence. Imaging and analysis of slides was performed at the University of Rennes. CRediT taxonomy: Conceptualization (EAM, GB), Data curation (EAM, GB, CC), Formal Analysis (GB, CC), funding acquisition (EAM, GB), Investigation (EAM, GB, CC), Methodology (EAM, GB), Project administration (EAM, GB), Resources (EAM, GB), Software (CC, GB), Supervision (EAM, GB), Validation (EAM, GB), Visualization (GB, CC), Writing original and review (EAM, GB, CC). All authors have approved the final version of the manuscript, agree to be accountable to all aspects of the work as presented, and all persons designated as authors qualify according to CRediT taxonomy.

## Acknowledgements

We thank S. Dutertre and X. Pinson at the Microscopy Rennes Imaging Center (MRic, *Biologie, Santé, Innovation Technologique* - BIOSIT, Rennes, France) for assistance. MRic is a member of the national infrastructure France-BioImaging, supported by the French National Research Agency (ANR-10-INBS-04). We are also grateful to R. L. Scherer for valuable preliminary experiments. Initial microscopy was carried out in the Microscopy Laboratory, Oklahoma State University, which received funds for purchasing the equipment from the NSF MRI award number 1919805.

## Funding

FACE Foundation: Transatlantic Research Partnership (EAM, GB), NIH NICHD R15 1R15HD105231-01 (EAM), French Center for Scientific Research/CNRS (GB), *Ligue Nationale Contre le Cancer* PhD fellowship IP/SC – 17653 (CC).

**IACUC approval number**: 20-07

